# Lack of Vector Competence in UK *Culex pipiens molestus* for Oropouche Virus

**DOI:** 10.1101/2025.05.07.652619

**Authors:** Jack Pilgrim, Victoria E Sy, Quentin Multeau, Thamil Vaani Komarasamy, Alain Kohl, Marcus SC Blagrove, Jolanta Tanianis-Hughes, Jolyon Medlock, Christopher Sanders, Matthew Baylis, Isabelle Dietrich

## Abstract

Oropouche virus (OROV) is an orthobunyavirus (*Peribunyaviridae*) that has caused recurrent outbreaks in South America and has recently expanded into the Caribbean, with various biting midge and mosquito species considered vectors. Recent imported cases to Europe and North America have raised concerns about the potential for local transmission in non-endemic areas. To assess this risk in the United Kingdom, we investigated the vector competence of *Culex pipiens molestus* (*Cx. molestus*), a human-biting mosquito common in urban environments. Laboratory-reared adult females were fed a bloodmeal containing a Cuban 2024 OROV outbreak strain (240023) and maintained at 27°C. Of 64 individuals tested at 12-s14 days post-infection via plaque assay, none were positive for OROV in bodies, indicating no evidence of infection and subsequently limited or no vector competence potential. These results provide evidence that UK populations of *Cx. molestus* are unlikely to support OROV transmission, thereby refining assessments of OROV emergence risk in temperate settings. Further studies are needed to test other putative UK mosquito vectors, as well as *Culicoides* biting midge species to fully assess the potential for OROV transmission in this region.

## Introduction

Oropouche virus (OROV) is an emerging orthobunyavirus (*Peribunyaviridae*) endemic to South America which has, periodically, caused epidemics in people living in the Amazon region. Since 2023, the largest outbreak on record has affected several countries in South America [1,2] and has also spread for the first time to Cuba [3], thus expanding its geographical range. Travellers, mostly from Cuba, have brought OROV to North America [4] and Europe [5], raising the risk of its spread and emergence to new areas.

OROV is transmitted in South America primarily by a species of Ceratopogonid biting midge, *Culicoides paraensis* [6], which is a diurnally active species that readily feeds on humans in urban settings. Other species of human-feeding biting midges may also be implicated (for example, *C. furens* and *C. insignis*), as are several species of mosquito, most notably the very widespread species in tropical climates *Culex quinquefasciatus*, which has frequently been found positive for OROV in the field and shown to be susceptible in the laboratory [7,8]. However, studies have shown that *Cx. quinquefasciatus* is not consistently competent following oral infection, indicating that evidence for *Culex* spp. as a vector is likely context-dependent [9]. Notably, recent genomic analyses suggest that the virus involved in the current outbreak is a reassortant [10], potentially exhibiting altered pathogenicity and an expanded range of competent vector species.

The introduction of OROV into other parts of the world by travellers from South America and the Caribbean raises the general question of whether midges and mosquitoes found in those regions are competent to transmit the virus to humans. Here, we consider the specific question of whether UK mosquitoes are competent to transmit OROV. *Cx. quinquefasciatus* is absent from the UK but it is a member of the *Cx. pipiens* assemblage, which includes the species *Cx. pipiens* sensu strictu. *Cx. pipiens* is found throughout the UK, often in close proximity to people. It occurs in the UK in three biotypes: the bird-biting *Cx. pipiens pipiens*, the human-biting *Cx. pipiens molestus* (hereafter *Cx. molestus*), and hybrids between the two [11]. All three biotypes feed on humans to differing extents and hence could present a public health threat from OROV. The highest risk vector is most likely *Cx. molestus*, which, where it is locally abundant [12], could feed on a traveller, become infected and transmit the infection to other people.

Here we investigated the competence of a colony of *Cx. molestus* established from collections in Surrey, southeast England, UK, for OROV. This was achieved by feeding adult females a virus-spiked bloodmeal, maintaining them at constant temperature and then, after several days, testing their bodies, legs and wings, and saliva for OROV by plaque assay as a measure of virus infection, dissemination and transmission, respectively.

## Materials and methods

### Mosquito maintenance

The *Cx. molestus* colony was derived from a mixed field population of *Cx. pipiens* s.l. (“Brookwood”) collected in 2011 from Surrey, UK [13]. Briefly, *Cx. molestus* were bred by rearing egg rafts from the mixed population, identifying emerging adults by biotype using molecular markers from pupal exuviae, and subsequently maintaining individual *Cx. molestus* biotypes through subsequent generations [14]. Egg rafts were hatched in 35 × 25 × 5 cm larval trays with 2 litres of tap water. Larvae were provided with 125 mg Brewer’s Yeast (Holland’s and Barrett, UK) added every 2 days until pupation. Adults then emerged in 30 × 30 × 30 cm BugDorm cages (BugDorm, Taichung, Taiwan) and were maintained at 27°C and 60% relative humidity in a 12:12 inverted light:dark cycle with 10 % sucrose solution and water until infection experiments.

### Virus

The 240023 strain of OROV, originally isolated from a febrile patient from Cuba [15] was kindly provided by the World Reference Centre for Emerging Viruses and Arboviruses (WRCEVA) at the University of Texas Medical Branch at Galveston, Texas, USA. Virus was passaged once on Vero cells (ATCC, Manassas, VA, USA) at low MOI for 5 days at 33°C and 5% CO2. To increase viral stock titres for mosquito infections, virus was concentrated over Amicon 100 kDa MW 15 ml columns (Merck, Feltham, UK). Virus stocks were titrated by plaque assay on Vero cells. Cells were incubated for 3 days at 37°C and 5% CO2 using Minimal Essential Medium (MEM; Fisher Scientific, Loughborough, UK) supplemented with 2% foetal calf serum (FCS; Fisher Scientific) and 0.6% Avicel (FMC Corporation/Roquette, London, UK) as overlay, fixed using 4% formaldehyde and stained using 0.1% (w/v) toluidine blue. All infection work was undertaken in specialised, licensed category 3 containment laboratories at The Pirbright Institute by trained personnel.

### Mosquito infection studies

Prior to infection, 9-20 day old female mosquitoes were starved of sucrose and water for 16 hours. They were cold-anesthetised and sorted into 4 oz espresso cups covered by fine mesh. On the day of infection, an infectious blood meal was prepared containing two-thirds defibrinated horse blood (TCS Biosciences, Buckingham, UK), one third virus at a final titre of 6.5 log10 PFU/ml and dATP (Thermo Fisher Scientific, Hemel Hempstead, UK) as a phagostimulant at a final concentration of 5 mM.

Mosquitoes were fed using a Hemotek feeding system (Hemotek, Blackburn, UK) set up at 36-37°C. Reservoirs were covered with one layer of hog gut and one layer of parafilm, and blood was offered to mosquitoes for 1-2 hours in a class III microbiological safety cabinet in the dark. Mosquitoes were cold-anesthetised and engorged individuals sorted into fresh cups. Non-fed females were discarded. Mosquitoes were maintained for 12-14 days at 27°C. 10% sucrose solution on cotton was provided throughout as source of nutrition and water.

### Mosquito sampling

On days 12-14 post blood-meal, mosquitoes were cold-anesthetised, legs and wings removed and stored in 200 μl virus isolation media (VIM). VIM consisted of Dulbecco’s Modified Eagle Medium (DMEM) supplemented with 2% FCS, 100 I.U./ml penicillin, 100 μg/ml streptomycin, 50 μg/ml gentamycin and 2.5 μg/ml amphotericin B (all Fisher Scientific). Saliva was collected via the capillary method using 200 μl pipette tips containing 20 μl of 50% sucrose (w/v) (Fisher Scientific)– FCS (1:1) for 30-60 minutes. Saliva was combined with 180 μl VIM. Bodies were then collected in separate tubes containing 200 μl VIM. Body, leg and wing sample tubes contained a 2.5 mm sterile stainless steel ball bearing (Dejay Distribution, Launceston, UK). Samples were stored at -80°C until further use.

### Testing of mosquitoes for OROV

Body samples were homogenised at 25 Hz for 2 × 40 seconds using a Tissuelyser II system (Qiagen, Manchester, UK). Homogenate was clarified by centrifugation at 10,000 xg for 5 minutes at 4°C and supernatant was used for serial titration. Viral titres were quantified by plaque assay as described above to determine infectivity in bodies. As no bodies positive for virus infection were found, leg and wing and saliva samples were not titrated in this study.

## Results

None of the mosquito bodies tested positive for OROV by titration (Table 1). The absence of OROV infection in mosquito bodies suggests that the virus is not competent to establish midgut infections in this species. As a result, further testing for viral dissemination or transmission via saliva or legs/wings was unnecessary for these individuals. Using the Clopper–Pearson Exact method, the upper bounds of the 95% and 99% confidence intervals for the true infection rate were estimated to be 5.60% and 7.95%, respectively.

**Table 1.** Infection outcomes in *Culex pipiens molestus* following oral exposure to OROV strain 240023. N/A = Not Applicable.

| Input (log<sub>10</sub> PFU/ml) | No. infected (no. tested)<br>[Infection rate 95% CI] | No.<br>disseminated | No.<br>transmitted |
| --- | --- | --- | --- |
| 6.5 | 0 (64) [0-5.6%] | N/A | N/A |

## Discussion

Our results could not find any evidence for vector competence of UK *Cx. molestus* for OROV, an outcome that is reassuring as *Cx. molestus* is a voracious human feeder that is closely related to a known vector, *Cx. quinquefasciatus. Cx. molestus* has a very focal distribution, mostly found underground and occasionally above ground [12]. Far more widespread and common in the UK is the *Cx. pipiens* biotype which is found throughout the country in both rural and urban environments [16]. It often breeds in rain harvesters or other water containers in gardens; it is probably the mosquito most frequently encountered by people and it will occasionally feed on them. However, while *Cx. molestus* is primarily a human feeder, *Cx. pipiens* is primarily a bird feeder. Recent assessment of vector competency for OROV of *Cx. pipiens* from the USA demonstrated one saliva positive out of 50 tested for the same virus used in this study (240023) suggesting that although transmission is possible, risk remains very low [15]. Given the close relatedness of *Cx. molestus* and *Cx. pipiens*, this suggests that vector competence for OROV across the species complex is likely to be minimal.

After feeding, the *Cx. molestus* in our study were maintained at 27°C until harvesting, mimicking achievable summer temperatures in England. Many mosquito-virus systems show increased infection and transmission rates at higher temperatures, despite local cooler environmental conditions [17,18]. However, this is not universally observed, with UK-derived *Cx. pipiens* showing high competence for the flavivirus Japanese encephalitis virus (JEV) when maintained at 18°C [19], suggesting that cold-adapted mosquitoes may, in some cases, be more susceptible at lower temperatures [20]. It is therefore important to follow up our initial experiments with studies at lower temperatures.

Similarly, there are several other UK species of mosquito that feed readily on people; for example, *Aedes vexans* and the salt marsh mosquito, *Ochlerotatus (Aedes) detritus*. The latter species has been shown to be competent to transmit several flaviviruses and alphaviruses (JEV, West Nile virus, Zika virus, Ross River virus) [19–22] and transmit the bunyavirus Rift Valley fever virus [23], while attempts to infect with dengue virus (a flavivirus) and chikungunya virus (an alphavirus) were unsuccessful [20]. Given its demonstrated competence for some arboviruses, investigation of the competence of *Och. detritus* for OROV should be prioritised.

A greater risk to people in the UK may arise from *Culicoides* biting midges which are diverse and widespread. The majority of species feed primarily on animals but some more generalist feeders are known to feed occasionally on people. This includes the widespread *C. obsoletus* species group and the Scottish biting midge, *C. impunctatus*, which reaches extremely high abundances in suitable habitats in the UK, especially the West of Scotland [24]. *C. impunctatus* is autogenous (meaning that mated females can lay their first egg batch without having taken a bloodmeal) and these high abundances can therefore be achieved where there are few or no suitable hosts. Consequently, the vast majority of adult *C. impunctatus* probably never take a bloodmeal. As transmission of OROV in the UK would require a vector to feed at least twice on humans (as there are no known reservoir species), in sparsely populated areas like Western Scotland it seems inherently unlikely that any *C. impunctatus* would achieve this, although there may be certain areas frequented by people where the probability of sequential human feeds is higher. Overall, the capacity of *C. impunctatus* to spread OROV is likely to be low, even if it is competent. Some *Culicoides* species (e.g., *C. obsoletus, C. pulicaris, C. punctatus*) can be a biting nuisance in peri-domestic environments [25] and investigation of their competence for OROV should be prioritised.

## Author contributions

JM, MB, MSCB, AK, CS and ID secured funding for this project. ID, MB, MSCB, JM, JP, CS and VES contributed to the conceptual development of the project. JP, JTH and QM maintained mosquitoes and prepared adults for infection experiments. ID and TVK prepared virus stocks and performed virus titrations. ID and VS conducted mosquito infection work. ID, MB and JP interpreted data and produced the first draft of the manuscript. All authors assisted in critical revision of the manuscript.

## Funding

This research was funded by a HPRU-EZI award to JM, MB, MSCB, JP, AK, CS and ID. Research conducted at The Pirbright Institute was further supported by funding from the Biotechnology and Biological Sciences Research Council (BBS/E/PI/230002C, BBS/E/PI/23NB0004, BBS/E/PI/23NB0003).

## Conflicts of interest

The authors declare that there are no conflicts of interest.

## Notes

### Competing Interest Statement

The authors have declared no competing interest.

## References

[1] Wesselmann KM, Postigo-Hidalgo I, Pezzi L, de Oliveira-Filho EF, Fischer C, de Lamballerie X, et al. Emergence of Oropouche fever in Latin America: a narrative review. Lancet Infect Dis 2024;24:e439–52. 10.1016/S1473-3099(23)00740-5.

[2] Sah R, Srivastava S, Kumar S, Golmei P, Rahaman SA, Mehta R, et al. Oropouche fever outbreak in Brazil: an emerging concern in Latin America. Lancet Microbe 2024;5:100904. 10.1016/S2666-5247(24)00136-8.

[3] Benitez AJ, Alvarez M, Perez L, Gravier R, Serrano S, Hernandez DM, et al. Oropouche Fever, Cuba, May 2024. Emerg Infect Dis 2024;30:2155–9. 10.3201/eid3010.240900.

[4] Morrison A, White JL, Hughes HR, Guagliardo SAJ, Velez JO, Fitzpatrick KA, et al. Oropouche Virus Disease Among U.S. Travelers - United States, 2024. MMWR Morb Mortal Wkly Rep 2024;73:769–73. 10.15585/mmwr.mm7335e1.

[5] Castilletti C, Mori A, Matucci A, Ronzoni N, Van Duffel L, Rossini G, et al. Oropouche fever cases diagnosed in Italy in two epidemiologically non-related travellers from Cuba, late May to early June 2024. Euro Surveill 2024;29. 10.2807/1560-7917.ES.2024.29.26.2400362.

[6] Pinheiro FP, Travassos da Rosa AP, Gomes ML, LeDuc JW, Hoch AL. Transmission of Oropouche virus from man to hamster by the midge Culicoides paraensis. Science 1982;215:1251–3. 10.1126/science.6800036.

[7] McGregor BL, Connelly CR, Kenney JL. Infection, Dissemination, and Transmission Potential of North American Culex quinquefasciatus, Culex tarsalis, and Culicoides sonorensis for Oropouche Virus. Viruses 2021;13. 10.3390/v13020226.

[8] Feitoza LHM, Gasparelo NWF, Meireles ACA, Rios FGF, Teixeira KS, da Silva MS, et al. Integrated surveillance for Oropouche Virus: Molecular evidence of potential urban vectors during an outbreak in the Brazilian Amazon. Acta Trop 2025;261:107487. 10.1016/j.actatropica.2024.107487.

[9] de Mendonça SF, Rocha MN, Ferreira F V, Leite THJF, Amadou SCG, Sucupira PHF, et al. Evaluation of Aedes aegypti, Aedes albopictus, and Culex quinquefasciatus Mosquitoes Competence to Oropouche virus Infection. Viruses 2021;13. 10.3390/v13050755.

[10] de Melo Iani FC, Pereira FM, de Oliveira EC, Rodrigues JTN, Machado MH, Fonseca V, et al. Travel-associated international spread of Oropouche virus beyond the Amazon. J Travel Med 2025;32. 10.1093/jtm/taaf018.

[11] Haba Y, McBride L. Origin and status of Culex pipiens mosquito ecotypes. Current Biology 2022;32:R237–46. 10.1016/j.cub.2022.01.062.

[12] Johnston CJ, Vaux AGC, Cull B, Medlock JM. Passive surveillance records including nuisance or suspected invasive/non-native mosquitoes in the United Kingdom, 2005-2021. J Eur Mosq Control Assoc 2023;41:35–46. 10.52004/JEMCA2022.0006.

[13] Manley R, Harrup LE, Veronesi E, Stubbins F, Stoner J, Gubbins S, et al. Testing of UK Populations of Culex pipiens L. for Schmallenberg Virus Vector Competence and Their Colonization. PLoS One 2015;10:e0134453. 10.1371/journal.pone.0134453.

[14] Jones L. Investigating the influence of hybridisation and larval habitat on the expression of life history traits of Culex pipiens in the United Kingdom. London School of Hygiene & Tropical Medicine, 2022.

[15] Payne AF, Stout J, Dumoulin P, Locksmith T, Heberlein LA, Mitchell M, et al. Lack of Competence of US Mosquito Species for Circulating Oropouche Virus. Emerg Infect Dis 2025;31:619–21. 10.3201/eid3103.241886.

[16] Widlake E, Wilson R, Pilgrim J, Vaux AGC, Tanianis-Hughes J, Haziqah-Rashid A, et al. Spatial distribution of Culex mosquitoes across England and Wales, July 2023 2025. 10.1101/2025.02.28.25322982.

[17] Jansen S, Heitmann A, Uusitalo R, Korhonen EM, Lühken R, Kliemke K, et al. Vector Competence of Northern European Culex pipiens Biotype pipiens and Culex torrentium to West Nile Virus and Sindbis Virus. Viruses 2023;15. 10.3390/v15030592.

[18] Jansen S, Heitmann A, Lühken R, Leggewie M, Helms M, Badusche M, et al. Culex torrentium: A Potent Vector for the Transmission of West Nile Virus in Central Europe. Viruses 2019;11. 10.3390/v11060492.

[19] Chapman GE, Sherlock K, Hesson JC, Blagrove MSC, Lycett GJ, Archer D, et al. Laboratory transmission potential of British mosquitoes for equine arboviruses. Parasit Vectors 2020;13:413. 10.1186/s13071-020-04285-x.

[20] Blagrove MSC, Sherlock K, Chapman GE, Impoinvil DE, McCall PJ, Medlock JM, et al. Evaluation of the vector competence of a native UK mosquito Ochlerotatus detritus (Aedes detritus) for dengue, chikungunya and West Nile viruses. Parasit Vectors 2016;9:452. 10.1186/s13071-016-1739-3.

[21] Mackenzie-Impoinvil L, Impoinvil DE, Galbraith SE, Dillon RJ, Ranson H, Johnson N, et al. Evaluation of a temperate climate mosquito, Ochlerotatus detritus (=Aedes detritus), as a potential vector of Japanese encephalitis virus. Med Vet Entomol 2015;29:1–9. 10.1111/mve.12083.

[22] Blagrove MSC, Caminade C, Diggle PJ, Patterson EI, Sherlock K, Chapman GE, et al. Potential for Zika virus transmission by mosquitoes in temperate climates. Proceedings of the Royal Society B: Biological Sciences 2020;287:20200119. 10.1098/rspb.2020.0119.

[23] Lumley S, Hernández-Triana LM, Horton DL, Fernández de Marco MDM, Medlock JM, Hewson R, et al. Competence of mosquitoes native to the United Kingdom to support replication and transmission of Rift Valley fever virus. Parasit Vectors 2018;11:308. 10.1186/s13071-018-2884-7.

[24] Stuart AE, Evans A, Brooks C, Simpson MJ, Cloughley JB, MacIntosh DF, et al. The biting midge of the West Highlands: fifty years of research. Scott Med J 1996;41:143–6. 10.1177/003693309604100505.

[25] Carpenter S, Groschup MH, Garros C, Felippe-Bauer ML, Purse B V. Culicoides biting midges, arboviruses and public health in Europe. Antiviral Res 2013;100:102–13. 10.1016/j.antiviral.2013.07.020.

